# Scientific civility and academic performance

**DOI:** 10.1101/2023.01.26.525747

**Authors:** Emma Camacho, Quigly Dragotakes, Isabella Hartshorn, Arturo Casadevall, Daniel L Buccino

## Abstract

In modern science, interdisciplinary and collaborative research is encouraged among scientists to solve complex problems. However, when the time comes to measure an individual’s academic productivity, collaborative efforts are hard to conceptualize and quantify. In this study, we hypothesized that a social behavior coined “scientific civility”, which encompasses civility, collaboration, cooperation, or a combination of these, enhances an individual’s productivity influencing their academic performance. To facilitate recognition of this unique attribute within the scientific environment, we developed a new indicator: the *C* score. We examined publicly available data from 1000 academic scientists at the individual-level, focusing on their scholarly output and collaborative networks as a function of geographic distribution and time. Our findings strongly suggest that the *C* score gauges academic performance from an integral perspective based on a synergistic interaction between productivity and collaborative networks, prevailing over institutionally limited economic resources and minimizing inequalities related to the length of individual’s academic career, field of investigation, and gender.

**Author Summary:** The increased connectivity between fields and specialties of science is undeniable. We propose a new indicator, the *C* score, to assess the value of collaborative efforts and research output to a scientist’s academic performance. This indicator reflects collaborative and interdisciplinary efforts and provides a measure of “scientific civility” and teamwork. The *C* score may be used as a decision-making tool to track career advancement within the academic environment. Along with other indicators such as the *h* index, the *C* score supports a more integrative and holistic assessment of an individual’s academic performance.

## Introduction

Modern science is a complex, self-organizing, and evolving global network of interconnected individuals through distinct forms of knowledge such as research articles, books, patents, and software, within time and space [1]. Traditionally, use of bibliometric indicators to assess scientific productivity and academic performance has revolved around quantitative measures regarding number of publications [2, 3], citation counts [4], and impact factor of the journal where research work was published [5]. Scientific success has been historically associated with academic tenure, securing external funding, and recognition by prestigious awards (e.g., Nobel Laureate) [6] within a tremendously competitive environment.

During the past decades, the notion of “team science”, defined as collaborative efforts (interdisciplinary or multidisciplinary) between and among groups of scientists who work together to solve a particular scientific challenge, has shifted research to a new era of team-authored publications dominating over the solo-authored research work. Often, this approach tends to be limited in time (defined duration) and/or space (a building with large open labs) [7, 8]. The advantages of collaborative work are multiple, including effectively joining specialized knowledge that could prompt scientific breakthroughs [9], pool resources to access cutting-edge instrumentation, gain visibility in newer networks, and increase productivity and citations [10], to name a few. Despite these benefits, when the time comes to measure academic productivity, promotions committees tend to overvalue the monadic scholar “genius” and potentially undervalue what appear to be overly collaborative or group enterprises [11]. Furthermore, multi-authored work has raised a relevant concern regarding fare credit allocation to coauthors, an aspect disregarded by the most used indicator of scientific achievement: the *h*-index [12]. The *h* index is a cumulative citation-based standard metric for quantifying the research output and impact of a scientist’s published work.

Regarding highly prolific collaborative work within our modern scientific enterprise, scientists’ social norms, notably civility, is the essential element that keeps the endeavor together [13]. Civility is more than being polite, it is the intentional choice of seeking common ground, disagreeing without disrespect, and genuinely caring about other people’s needs and beliefs [14]. The core constituents of civility – relational competence and purposeful poise [15]– are essential for productivity and morale where there may be individuals with four to five decades of age difference working together, as well as people of different races, nationalities, gender identities, sexual orientations, and many other sometimes competing equity interests. The American workplace [16] is among the most diverse in the world and the academic milieu is no different. Whether in a laboratory, a clinic, or a classroom, successful scientists in academia must be able to operate effectively in a diverse and dynamic environment, whether they be knowledge generators or disseminators, clinicians, educators, or administrators. Workplace civility has been defined as behaviors that help to preserve the norms of mutual respect and collaboration among co-workers [17, 18]. The co-inventors of the transistor and winners of the 1956 Nobel Prize in Physics, William Shockley, Walther Houser Brattain, and John Bardeen, provide a sad illustration of this case. Their brilliant years of collaborative work at the Bell Labs ended, in part to due to a clash of personalities and to Shockley’s advocacy for eugenics and racist views [19].

In this study we investigated the hypothesis that the academic performance and impact of an individual’s scientific output was strongly associated with a social behavior we called “scientific civility” to capture the constellation of attributes that contribute to successful human interactions in science. To quantify this observation, we developed an index termed the *C* score, where the C stands for *C*ivility, *C*ooperation, *C*ollaboration, or a combination of these, which for simplicity we shorthand to civility with the understanding that it encompasses the other C’s. We propose that the *C* score provides an integrative and holistic approach to assess an individual’s academic performance as the result of their successful scientific collaborations. This characteristic is an important attribute of scientific citizenship whose measure should be considered a valuable tool in professional and promotional decision-making scenarios.

## Results

### Development of a new metric: The *C* score

Using a binary scoring system, we operationalize the abstract construct of “scientific civility” into a measurable observation that ranges from 0 to 1 point. At a defined time of a scientist’s career, their *C* score is determined as *C* = *SOY* + *CN*, where SOY (Scholarly Output per Year) corresponded to academic productivity and CN (Collaborative Networks) corresponded to collaborative efforts over time and geographic distribution. SOY and CN are equally weighted variables that individually account for 50% of the total score.

In this study, academic productivity corresponded to published articles of individuals affiliated to research academic institutions. Therefore, the scholarly output refers to Research Output per Year (ROY) and equals to the total number of publications generated during a scientist’s academic career normalized by year. To calculate an individual’s *C* score, for the ROY component of the equation a point is given according to the level of productivity that matches the ROY ≥20, ≥15, ≥10, ≥5, and >0. To validate the accuracy of these levels of productivity, we assessed the ROY distribution among a sample of 928 scientists in academia, which included 744 Tenure-Track Professors and 103 Nobel Laureates (**Fig. 1**). The calculated median ROY was 4.8. Notably, independently of the categorization used (schools, academic ranks, and gender, **Fig 1A, Fig 1B**, and **Fig 1C**, respectively), we found a non-symmetric right skewed distribution demonstrating that the number of individuals decreased as the level of productivity increased. These results suggested that motivation for prolific scholarly productivity among scientists in academia is driven by individual intrinsic factors. Therefore, individuals displaying the highest productivity level were rewarded with a maximum of 5 points.

**Fig. 1.**
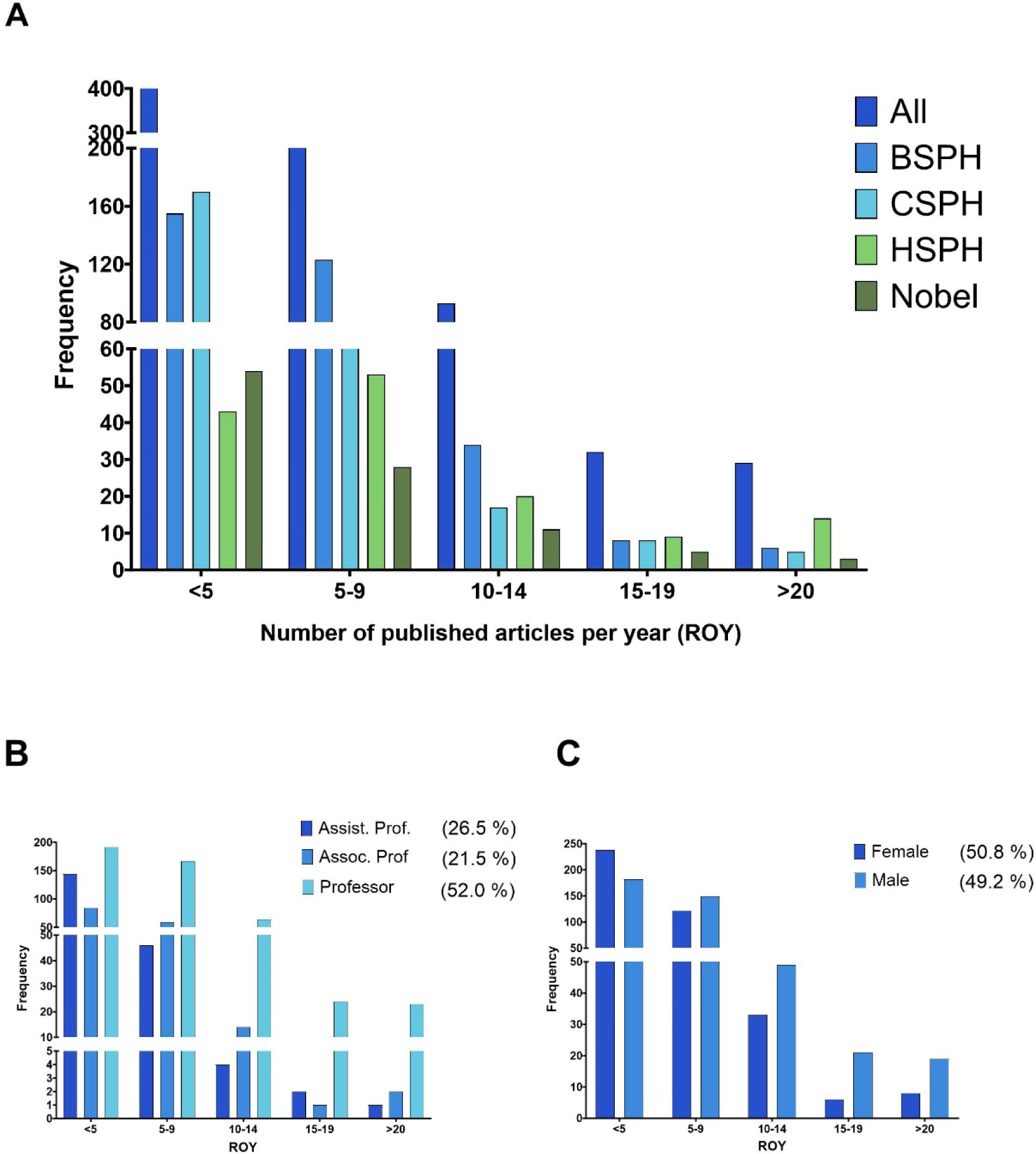
Prolific scholarly productivity among scientists in academia is intrinsically motivated. **A)** Distribution of research productivity based on the number of published articles per year, or research output per year, (ROY) among tenure-track professors from the Bloomberg Johns Hopkins University School of Public Health (BSPH), Columbia University Mailman School of Public Health (CSPH), Harvard T.H. Chan School of Public Health (HSPH), and Nobel Laureates in Chemistry, Physics, Economic Sciences, and Physiology from 2010 to 2020. **B)** Distribution of productivity from same population described in panel A but plotted according to their academic ranks: Assistant Professor (Assist. Prof), Associate Professor (Assoc. Prof), and Professor (includes Full Professors, Chairs, and JHU members of the National Academy of Science). **C)** Distribution of productivity from same population described in panel A but plotted according to their assigned gender.

For the CN component, we created sub-categories that measured levels of engagement as a function of geographic distribution and time (**Fig. 2**). These are the following:

1. Global collaboration network, accounts for the existence of collaborative projects with investigators affiliated with institutions from diverse cultural backgrounds using as reference the seven continents model: North America, South America, Europe, Asia, Africa, Oceania, and Antarctica. A point was given to an individual, if collaborative efforts with other scientists located in ≥4 continents have occurred during the length of their career.
2. Long-term collaborations sustained within the oldest 12 years or from the start of the academic career. For this study we used the time frame 2006-2017. This refers to the scientist’s ability to develop and maintain prolific productivity with one or more investigators over a decade. Points were assigned according to the level of productivity with unique investigators: Minimum of 30 published articles, Minimum of 15 published articles, and Minimum of 10 published articles. A maximum of 3 points could be accounted for this criterion.
3. Brand-new collaborations developed within the last 3 years (2018-2020). This refers to the scientist’s ability to intellectually engage with new investigators to develop, publish, and maintain collaborative projects that further expand their main research area and/or create a new line of investigation. As done previously, points were assigned according to the level of productivity with unique investigators: Minimum of 10 published articles, and Minimum of 5 published articles. A maximum of 2 points could be accounted for this criterion.

**Fig. 2.**
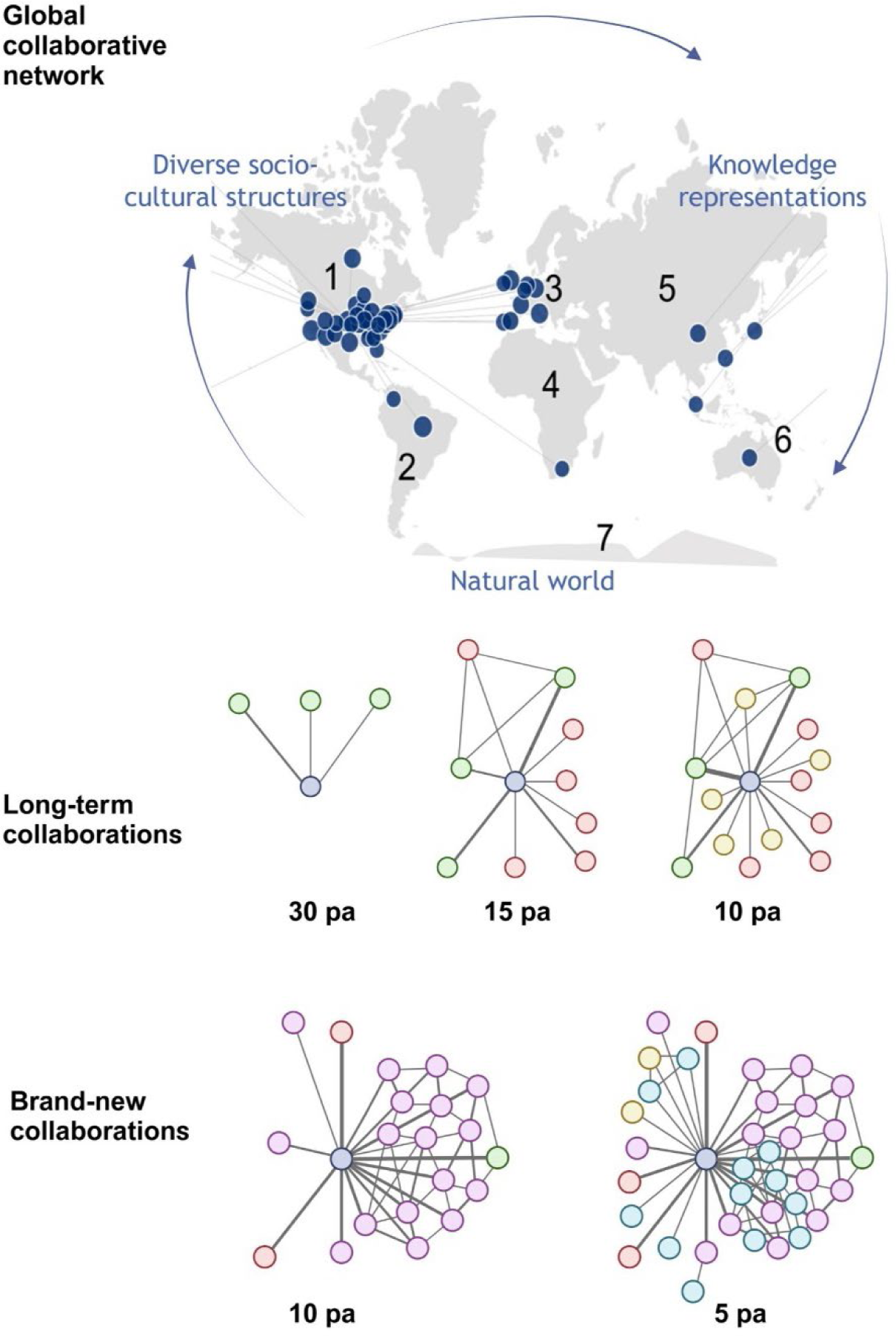
Collaborative networks (CN) built during a scientist academic career reflect their sphere of influence. Diagram of collaborative efforts assessed as a function of geographic distribution and time. *Upper panel*, **Global collaborative network** captures the ability to establish interactions (collaborative projects) among diverse socio-cultures structures (culturally diverse individuals) through knowledge representations (e.g. research articles, books, patents, software, or other scholarly output) and the natural world (research institutions across the seven continents) connected via formal or informal flows of information. *Middle panel*, **Long-term collaborations** capture the ability to develop and maintain prolific productivity with unique scientists over a decade. *Bottom panel*, **Brand-new collaborations** refer to the ability to intellectually engage with new investigators. Thickness of interconnecting lines represents level of engagement between individuals in terms of number of published articles (pa). Dark blue represents an individual’s network that is being assessed.

### The *C* score across research fields

We hypothesized that sustained collaborative networks are an important characteristic of successful academic/scientific performance. To validate or refute this hypothesis, we first set out to analyze the academic profiles of individuals from three top 5 Schools of Public Health (SPHs) within the United States of America (USA) [20], which embrace numerous departments with diverse nature of research approaches to study problems. These included tenure-track professors (i.e., Assistant Professor, Associate Professor, and Professor) with a primary affiliation to a department within their respective School of Public Health. A Spearman correlation (ρ) was computed to investigate whether there is a relationship between the *C* score and *h* index. The analysis revealed a strong, positive correlation between the two variables, which was statistically highly significant (ρ = 0.7640, *N* = 744, *P* < 0.0001). Hence, the *h* index was associated with the *C* score. Mean *C* score and mean *h* index were 0.43 and 39.15, respectively (**Fig.3A**). An individual analysis of each school exhibited significant differences between their median *C* score (Kruskal-Wallis test, 0.53; 0.43 and 0.35 for HSPH, BSPH, and CSPH, respectively) (**Fig 3B**). Furthermore, based on the type of lab environment, we grouped departments into dry, humid, or wet labs. We considered as dry labs those in the departments of: Biostatistics (BIO), Global health and Population (GHP), Health, Behavior and Society (HBS), Health Policy and Management (HPM), International Health (IH), Mental Health (MH), Population and Family Health (PFH), Population, Family and Reproductive Health (PFRH), Social and Behavioral Sciences (SBS), and Sociomedical Sciences (SS); as humid labs those in departments of: Environmental Health and Engineering (EHE), Environmental Health (ENV), Epidemiology (EPI), and Nutrition (NUT); and as wet labs those in departments of: Biochemistry and Molecular Biology (BMB), Immunology and Infectious Diseases (IID), and Molecular Microbiology and Immunology (MMI). We found that there was a strong correlation between these two variables, lowest ρ = 0.47, *P* = 0.0508 to highest ρ = 0.86, *P* < 0.0001 for A and G departments, respectively (**Fig. 3B**). Further distinction between the type of lab environment (dry, humid, or wet) demonstrated that only scientists associated to humid labs had a significantly higher median *C* score than those grouped as working in dry or wet labs (Kruskal-Wallis test, 0.45 *N* = 296; 0.37 *N* = 395; and 0.35 *N* = 53, respectively) (**Fig. 3C**). Altogether, these results suggest that the *C* score could be a potential tool to analyze in an integrated manner both scholarly productivity and collaborative efforts, independently of the research field.

**Fig. 3.**
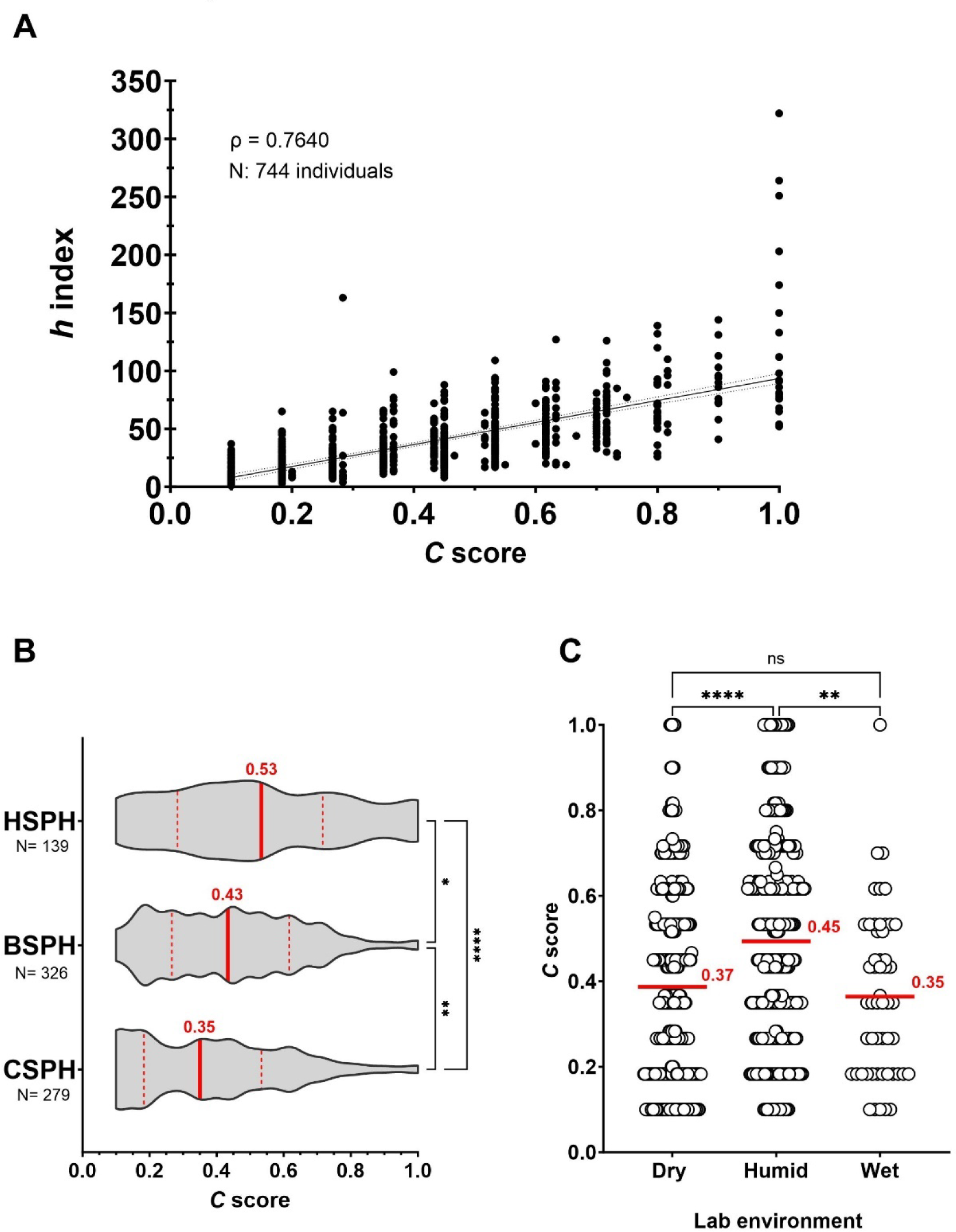
*C* score, a prospective metric to integrally analyze academic output and collaborative initiatives across diverse research domains. A scientist’s *h* index shows a positive strong relationship with the *C* score. **A)** Case study of three diverse SPHs in the USA: Harvard T.H. Chan School of Public Health (HSPH), Bloomberg Johns Hopkins University School of Public Health (BSPH), and Columbia University Mailman School of Public Health (CSPH). B) *C* score distribution in each SPH. C) *C* score distribution by grouping departments according to the type of laboratory environment.

### Impact of career length, academic ranks, and gender on the *C* score

Next, we tested whether the number of years in academia since a scientist published their first article would impact the *C* score. For this analysis, we also included other JHU professors not affiliated to the BSPH being recognized as members of the National Academy of Sciences (NAS) or holding leadership positions as Chair of Department/Center. These included 83 individuals from the School of Medicine, Whiting School of Engineering, and Krieger School of Arts and Sciences. The length of the academic career demonstrated no association with the *C* score, suggesting that academic performance of tenure-track professors is not tight to the duration of their career (ρ = - 0.0004, *N* = 827, *P <* 0.0001 (**Fig. 4A**). Nonetheless, we noted that when breaking down the analysis by tenure-track faculty ranks of Assistant Professor (Assist. Prof.), Associate Professor (Assoc. Prof.), and Professor (Prof.), median *C* score reflected progress within the academic career track with marked differences (Kruskal-Wallis test, 0.27 *N* = 197, 0.37 *N* = 160, and 0.52 *N* = 470, respectively) (**Fig 4B**). The overall analysis by gender exhibited significantly higher *C* score for male in comparison to female tenure-track professors (Kruskal-Wallis test, 0.45 *N* = 420 and 0.35 *N* = 404, respectively) (**Fig. 4C**). However, when comparing female and male professors with equal academic rank including Professor Chair, median *C* score revealed no gender differences (Kruskal-Wallis test, 0.27 *N* = 131, 0.27 *N* = 66; 0.36 *N* = 88, 0.43 *N* = 77; 0.48 *N* = 154, 0.53 *N* = 238, 0.27 *N* = 34, 0.45 *N* = 44, respectively) (**Fig. 4D**). Altogether, these results demonstrate the *C* score’s potential value for assessing career advancement without gender bias.

**Fig. 4.**
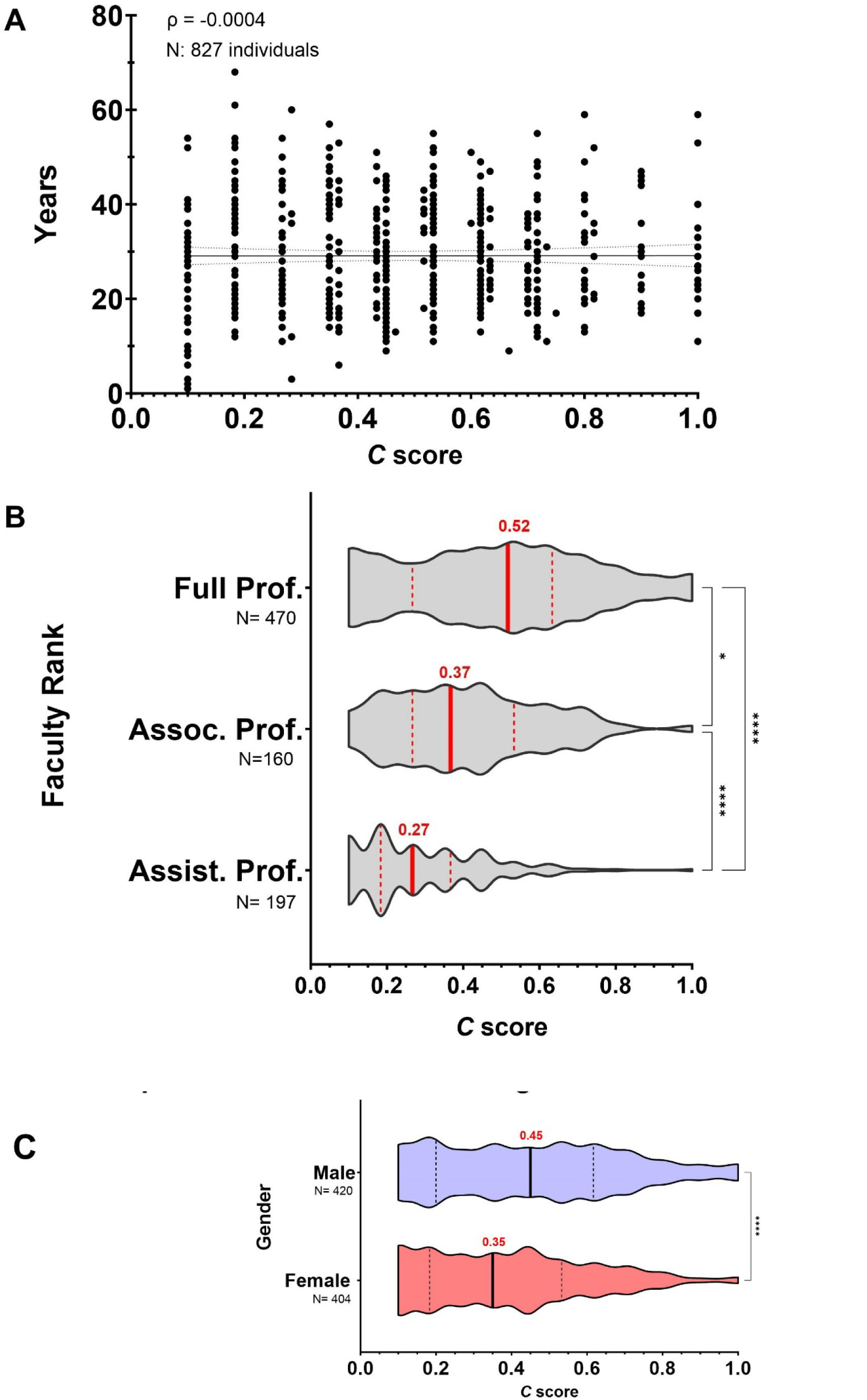
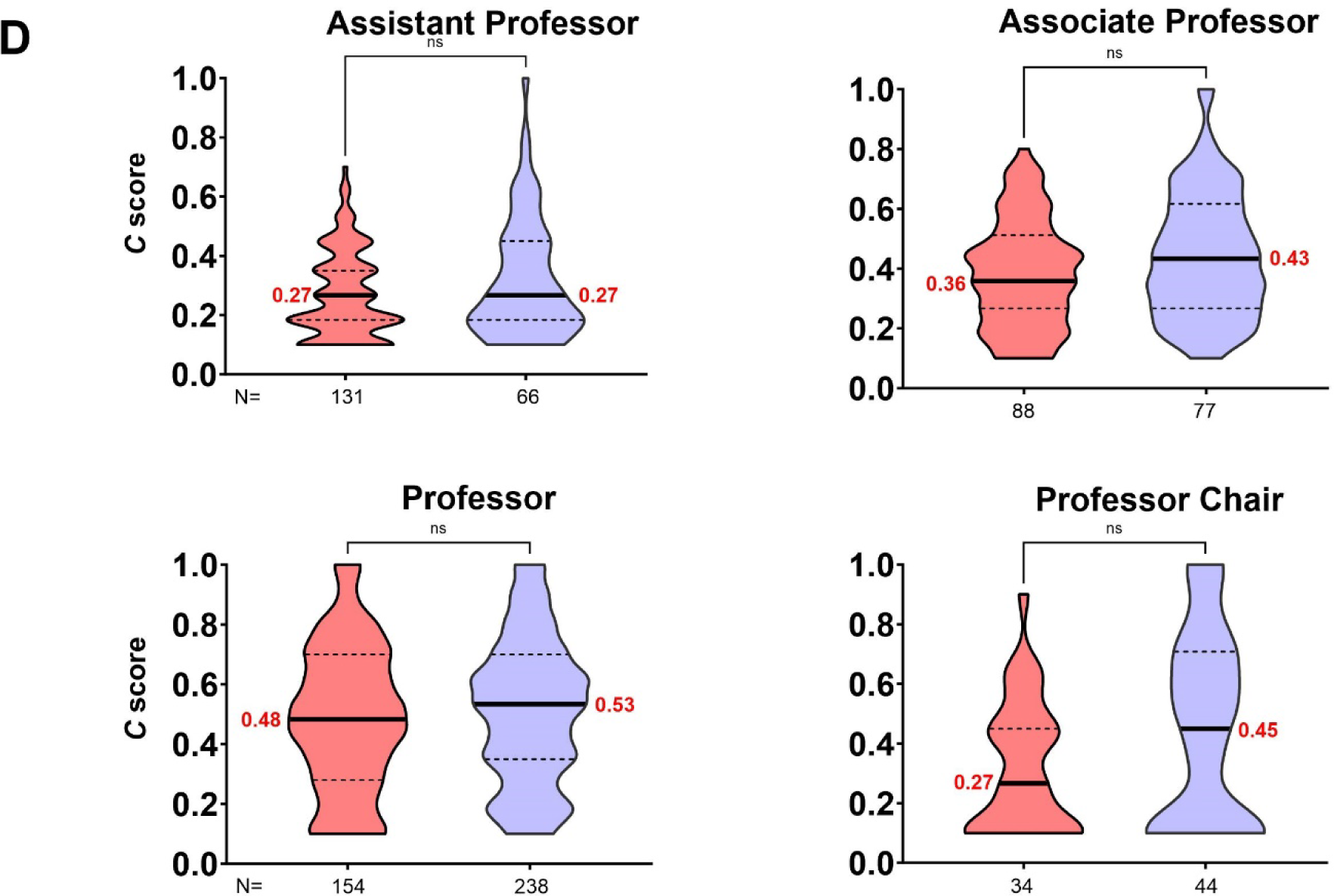
*C* score is a strong index to track advancement of an individual’s academic career revealing no gender bias between peers of equal rank. **A)** The *C* score is not affected by the length of an individual’s academic career. **B)** Median *C* score of academic ranks reflect progress within the academic career track. C) *C* score comparisons between male and female professors show strong differences. **D)** Gender disparities disappeared when comparing *C* score between peers of equal academic rank.

### Likelihood of recognition and dynamic behavior of the *C* score

We also investigated whether a high *C* score was associated with individuals whose significant scientific achievements were recognized by a Nobel Prize. For this purpose, we analyzed the profiles from 103 Nobel Laureates from 2010-2020 in the categories of Physiology, Chemistry, Physics, and Economic Sciences. Like the tenure-track professors, for Nobel Laureates the *h* index and *C* score were positively correlated, ρ = 0.7877, *P <* 0.0001 (**Fig. 5A**). Mean *C* score-*h* index pairings for this group were 0.47-77.44 while by individual categories were 0.43-91.75, 0.46-75.1, 0.18-39.17, and 0.48-95 for Chemistry, Physics, Economic Sciences, and Physiology, respectively **(Fig. 5B)**. A strong to very strong positive relationship between the two variables was found for all four categories, including those in the field of economics (ρ = 0.5087, *N* = 22, *P* = 0.0156). For both individual and shared Nobel Prizes, we found that the *C* score of awardees spanned the whole range of the index (from 0 to 1). This implies that Nobel recognition can occur at the productivity peak of an individual, long after peak productivity, or even when the scientist has only published a few articles and their discoveries were considered to confer the greatest benefit on mankind. Interestingly, we noted that throughout a scientist’s career the *C* score could exhibit a dynamic behavior. To illustrate this observation, we selected 8 Nobel Laureates from the same category whose career length was at least 50 years (**Fig. 5C**). Over the 50-year period of these individual’s scientific careers, the *C* score increased, decreased, or remained steady. These distinct patterns reflect diverse levels of engagement in collaborative projects.

**Fig. 5.**
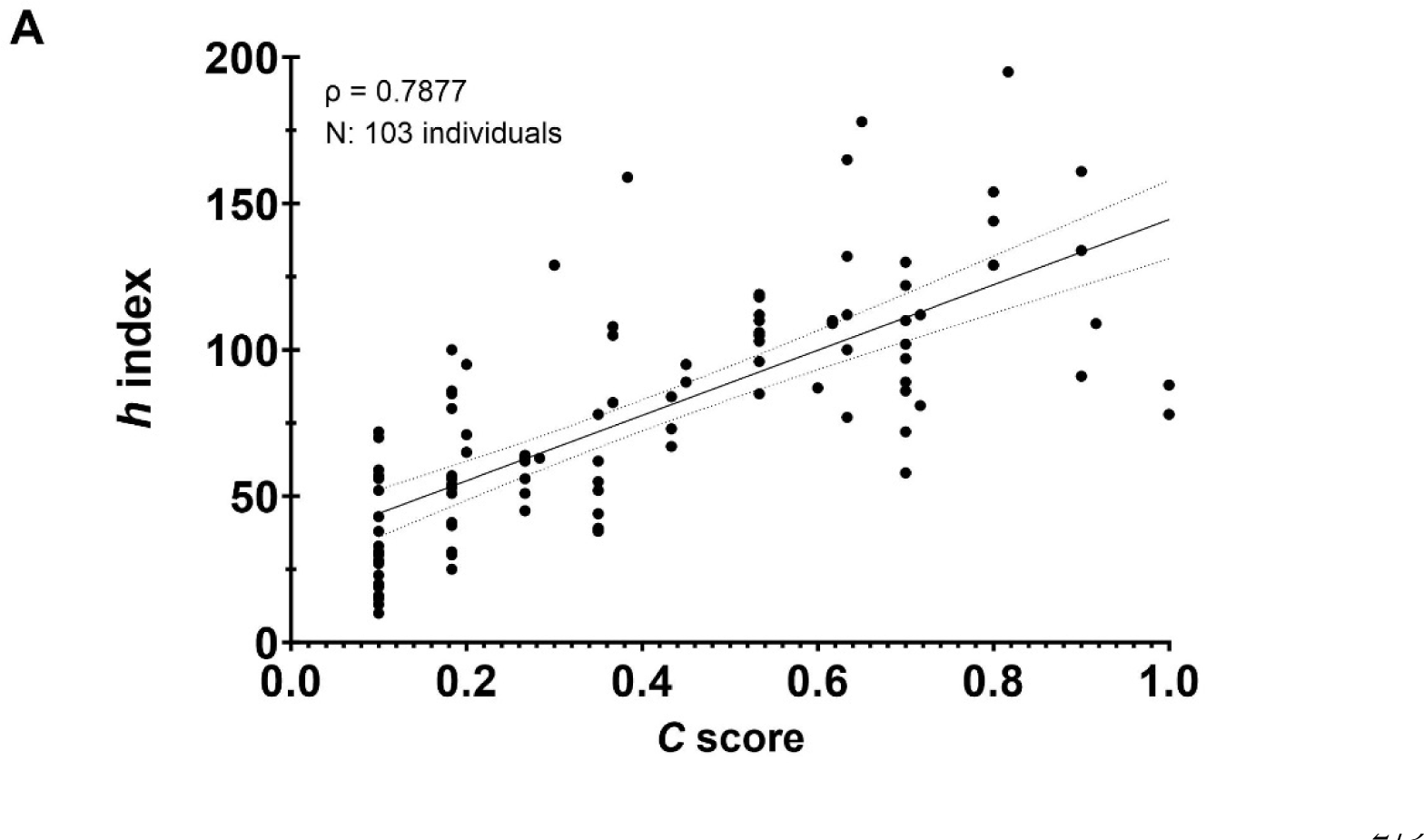
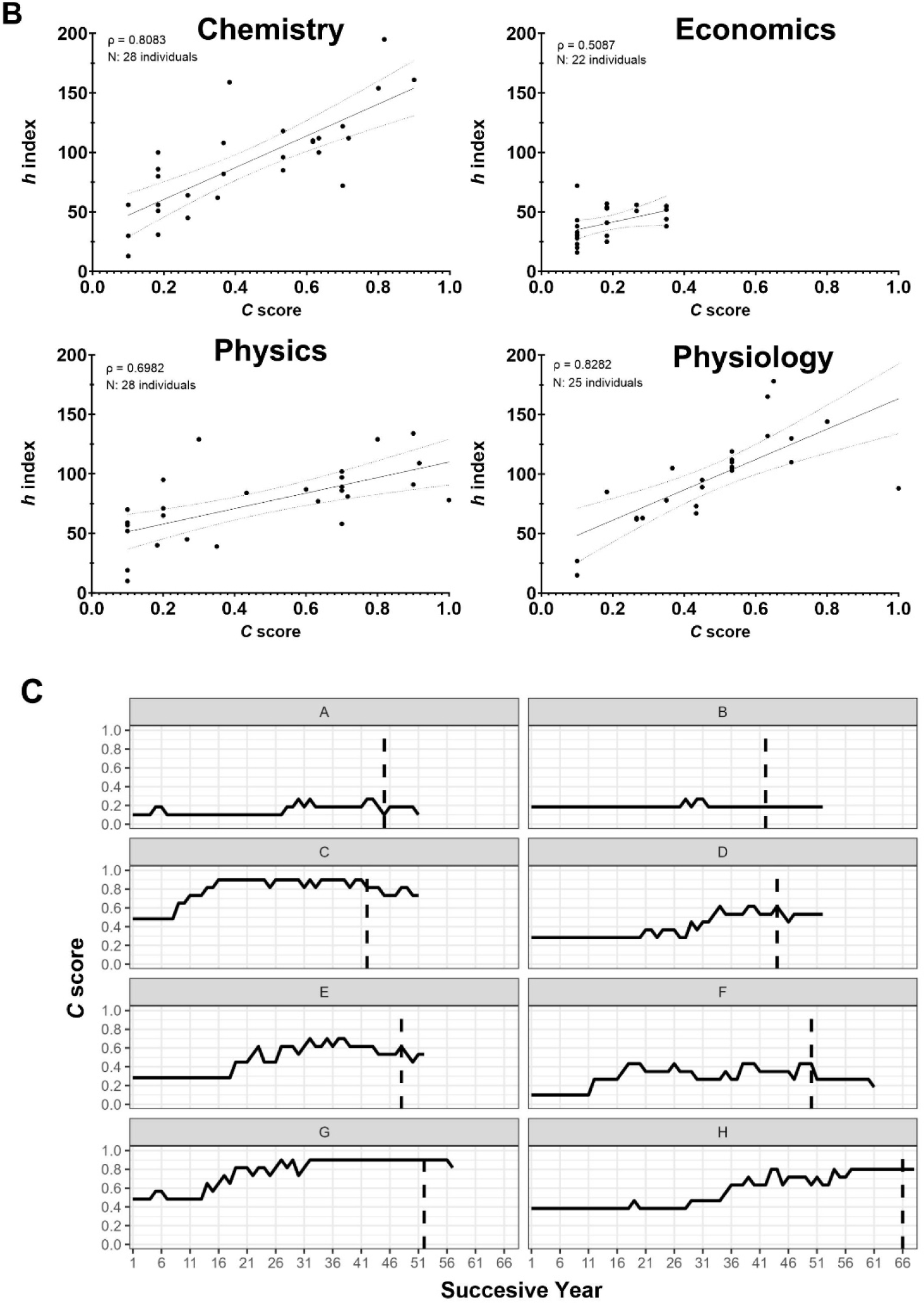
The *C* score reflects the likelihood of being recognized by a prestigious award and does not reflect a cumulative pattern. **A)** Case study of Nobel Laurates from 2010-2020 in four categories. **B)** Individual categories of Nobel Prizes also demonstrated a strong to very strong association between the two variables. **C)** The *C* score dynamics during the length of a scientist’s career reflect distinct levels of engagement with collaborative projects. The dotted line refers to the year that the selected scientists were awarded a Nobel prize.

### Economic environment and individual interests/values

Furthermore, since the productivity and relevance of a scientist’s research output is highly dependent on the economic resources of the institution to which he or she is affiliated [21], we assessed whether the impact and record of scientific achievements among researchers from countries with limited economic resources also associates with high *C* scores. Starting with the premise that the *C* score was useful to estimate the broad impact of a scientist’s published work from the biological sciences, we selected a group of individuals from outside the USA who were elected to the American Academy of Microbiology [22]. The Academy is the honorific leadership group within the American Society for Microbiology (ASM), one of the largest life science societies in the world [23]. To be elected for fellowship, two basic qualifications must be met: 1) Recognition at the national or international level, and 2) Outstanding and original contributions to the microbial sciences [22]. In addition, we considered culturally diverse individuals from nine countries defined as Middle-Income Countries (MICs) according to the World Bank (e.g., Argentina, Brazil, South Africa, India, Mexico, Lebanon, Russia, China, and Bangladesh). These countries are home to 75% of the world’s population and 62% of the world’s poor [24]. Like the groups previously analyzed, we found a strong positive correlation between the *C* score and *h* index, ρ = 0.8084, *N* = 37, *P <* 0.0001. (**Fig. 6A**). On another measure, we examined the *C* score from authors involved in retracted articles. Using the retraction watch database [25], we selected individuals who were affiliated with institutions in the USA and co-authored research articles retracted between 01/01/2015 to 12/31/2022 due to falsification/fabrication of image or paper mill. These scientists also revealed a strong association between *h* index and *C* score, ρ = 0.6912, *N* = 35, *P <* 0.0001. (**Fig. 6B**). We found no significant differences between the median *C* score of these two groups (Mann-Whitney test, 0.52 and 0.45, respectively) (**Fig. 6C**). These results are consistent with the notion that increased collaborative efforts across national and international borders can neutralize gaps in research productivity due to economic limitations [26]; however, a high *C* score cannot anticipate the motivation factors influencing collaborative work.

**Fig. 6.**
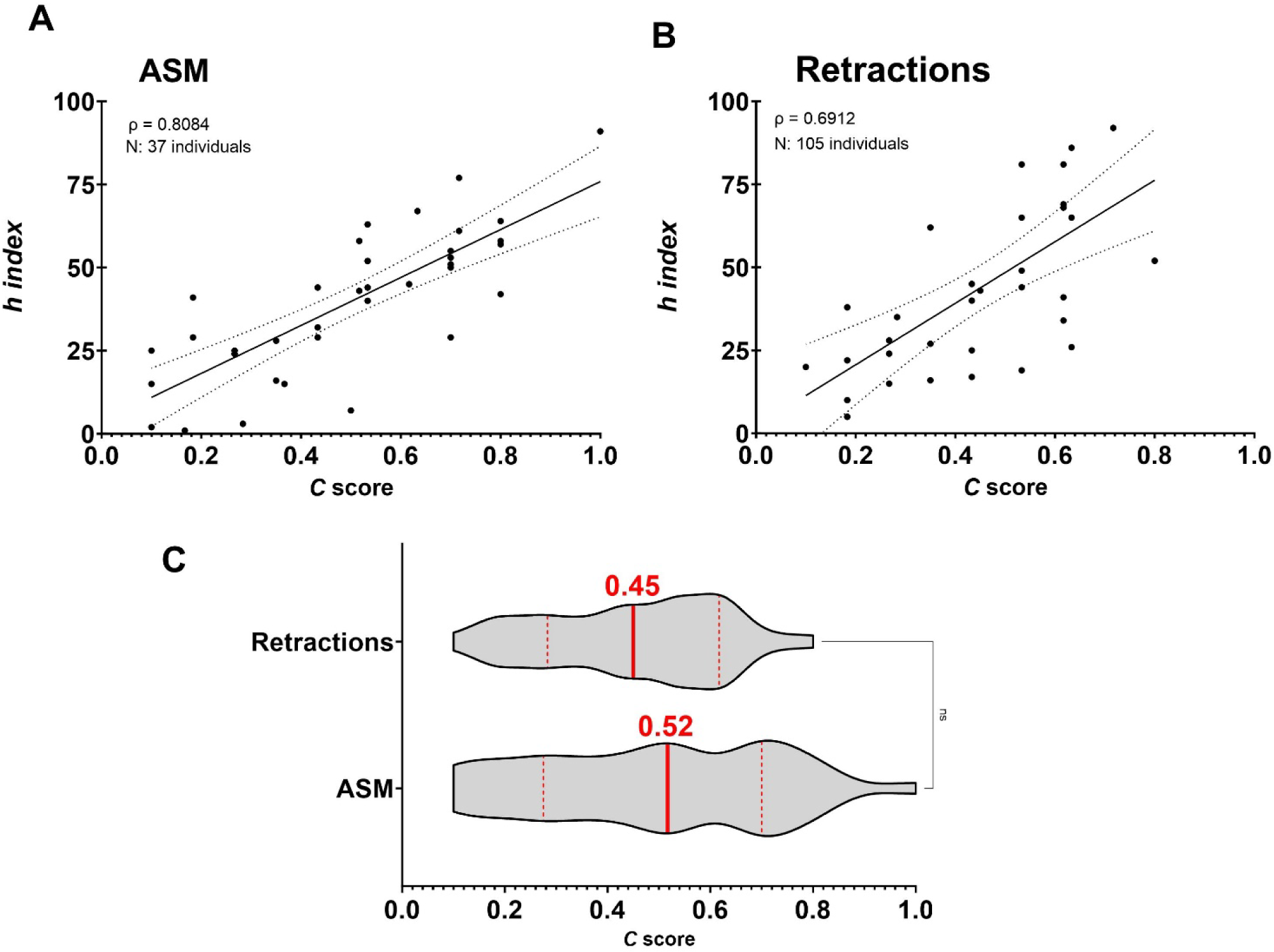
Active collaborative networks can overcome economic limitations but are unlikely to reflect an individual’s motivation for science. **A)** Case study of ASM International Fellows from Middle Income Economies countries shows that intense collaborative efforts serve as catalyst for increased productivity. **B)** Case study of retracted authors also reveals high collaborative work driving enhanced productivity. **C)** The *C* score does not portray the motivational factors impacting productivity.

### Sustained collaborative networks, Maslow’s hierarchy of motivational needs, and *C* score

Collaborations can have a large effect in science careers from more productivity and citations to greater visibility [1]. Therefore, we determined whether the *C* score correlates with unique names of collaborators with the prediction being that investigators with higher *C* value would have more unique names identified with them. As expected, we found a very strong and positive correlation between the total number of unique collaborators and the *C* score, (ρ = 0.7904, *N* = 827, *P <* 0.0001) (**Fig. 7A**). Given that the more publications people have the more likely they are to have unique collaborators associated with them, this strong association may not be indicative of large and sustained collaborative networks. Therefore, we limited the analysis to first and last authors considering that collaborative efforts between the lead and senior authors of a publication are more likely to suggest direct work between these individuals. This analysis indeed revealed a very strong correlation between the *C* score and the total number of unique first and last authors within a scientist’s publications (ρ = 0.7942, *N* = 827, *P <* 0.0001) (**Fig. 7B**). Next, relying on our data of the total number of unique first and last authors (Unique FL authors), we evaluated whether academic performance (*C* score) in tenure-track professors is rooted in human motivation. Based on Maslow’s hierarchy of needs as our conceptual framework [27, 28], we established three motivational levels by referencing the progressive pattern of median *C* scores that we captured throughout the tenure-track career (**Fig. 4C**). These levels were defined as follows: *C* scores below 0.3, between 0.3 and 0.8, and above 0.8 corresponded to *C-I*, *C-II*, and *C-III*, respectively. *C-I* depicts individuals driven by physiological, safety, and social needs, *C-II* are influenced by a deeper appreciation and understanding of their surroundings, and *C-III* portrays those individuals that seek to achieve their greatest potential and/or help others to achieve it (**Fig. 7C**). As motivational factors driving productivity move away from the physiological needs, collaborative networks exhibit highly significant growth (Tukey’s multiple comparison test, 27, 103, and 305 unique FL authors, respectively) (**Fig. 7D**). Likewise, productivity in terms of *h* index also demonstrated a similar distribution with highly significant differences between the three levels (Tukey’s multiple comparison test. 15, 43, and 90 h index for *C-I*, *C-II*, and *C-III*, respectively); nonetheless, given the accumulative nature of the *h* index, this metric is unlikely to reflect motivational factors driven productivity (**Supp** **Fig. 1**). These results support the notion that large and sustained collaborative networks catalyze scholarly outcomes contributing to a scientist’s academic performance.

**Fig. 7.**
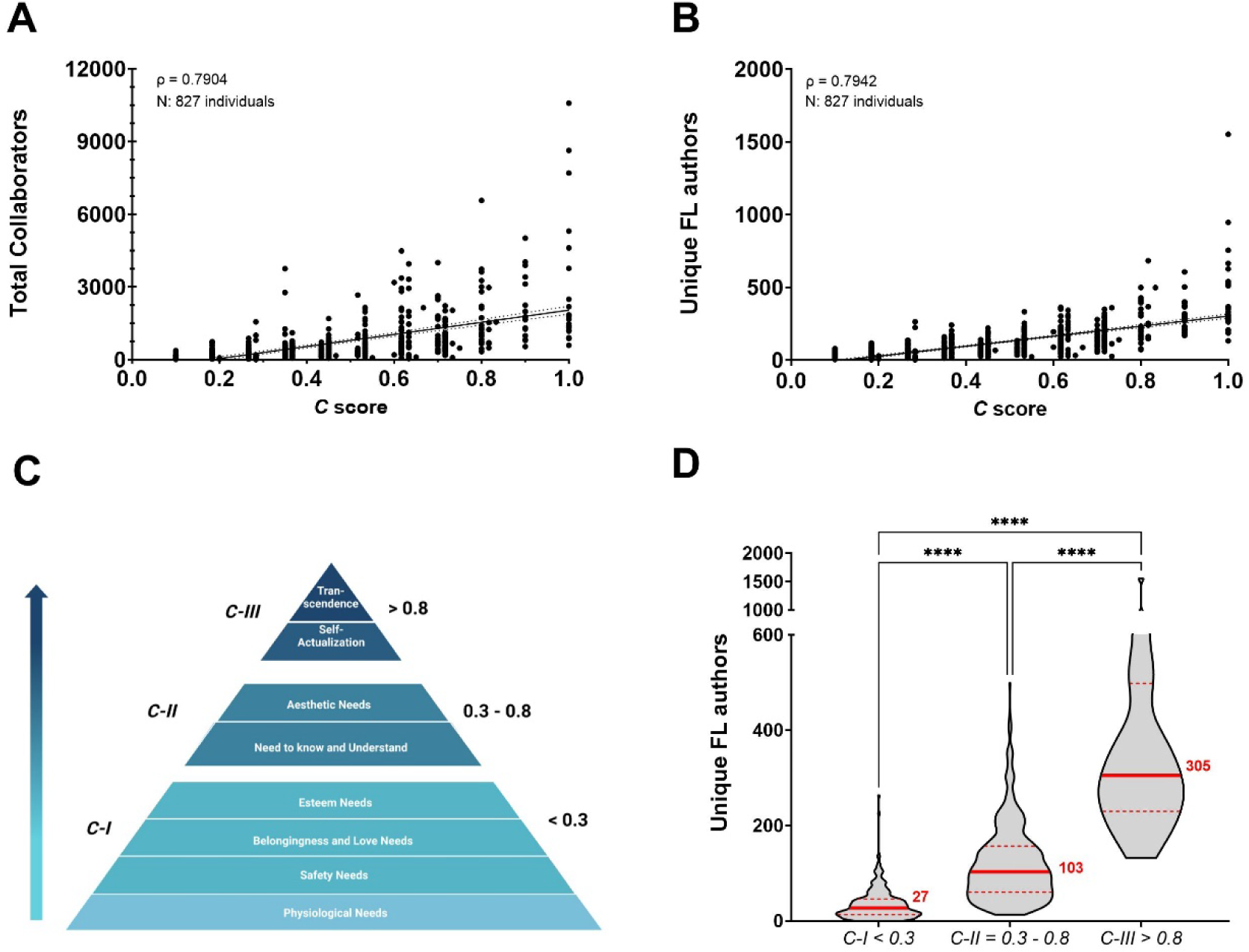
Large and sustained collaborative networks bolster an individual’s academic performance in response to distinct motivational needs. **A)** Total number of unique collaborators indicates a strong correlation with the *C* score. **B)** Number of unique First and Last author collaborators, indicative of efficient collaborative tendencies, reveal a very strong correlation with the *C* score. **C)** Using as conceptual framework for human motivation Maslow’s hierarchy of needs, *C* score levels portray the motivational factors impacting productivity. **D)** A distinct network size with highly significant increments is shown according to *C* score level.

### Collaborative networks, productivity, team size, sphere of influence, funding, gender disparities, and *C* level

We set to probe whether *C* score levels can reflect insights on the characteristics of research collaboration networks to achieve success in academia. The analysis of the size of collaborative networks between female and male scientists demonstrated than the median number of unique FL authors is significantly smaller for female than male scientists, independently of the *C* level (Mann-Whitney test, 23 vs 33; 83 vs 121; and 270 vs 367 for *C-I*, *C-II*, and *C-III*, respectively) (**Fig. 8A, upper panel**). Nonetheless, reduced extension of the collaborative network from female scientists did not seem to negatively affect their ROY in comparison to male (Mann-Whitney test, 2 vs 2; 6 vs 7; and 22 vs 20 for *C-I*, *C-II*, and *C-III*, respectively) (**Fig. 8A, lower panel)**. Next, to estimate a scientist’s team size we computed the median number of authors per article showing highly significant differences between the *C* levels (Kruskal-Wallis test, 3 *N* = 308, 5 *N* = 471, and 6 *N* = 47, for *C-I*, *C-II*, and *C-III* respectively) (**Fig. 8B**). Scientists within *C-I* and *C-II* levels tent to publish research articles comprising up to 21 and 31 co-authors, respectively, while *C-III* scientists tent to be associated to publications with a limited number of coauthors (up to 18) and no solo-authored work. In addition, an analysis of the sphere of influence of these individuals determined by the average number of countries with scientific connectivity showed a significantly increasing pattern as *C* level augmented (Kruskal-Wallis test, 11, 28, and 53 countries for *C-I*, *C-II*, and *C-III* level, respectively) **(Fig. 8C)**. These results demonstrate that unique differences on the extension of collaborative efforts between female and male scientists do not adversely impact female productivity; while both female and male research teams are composed of 3 to 6 individuals, their ability to connect with the world can expand to up to over 100 countries.

**Fig. 8.**
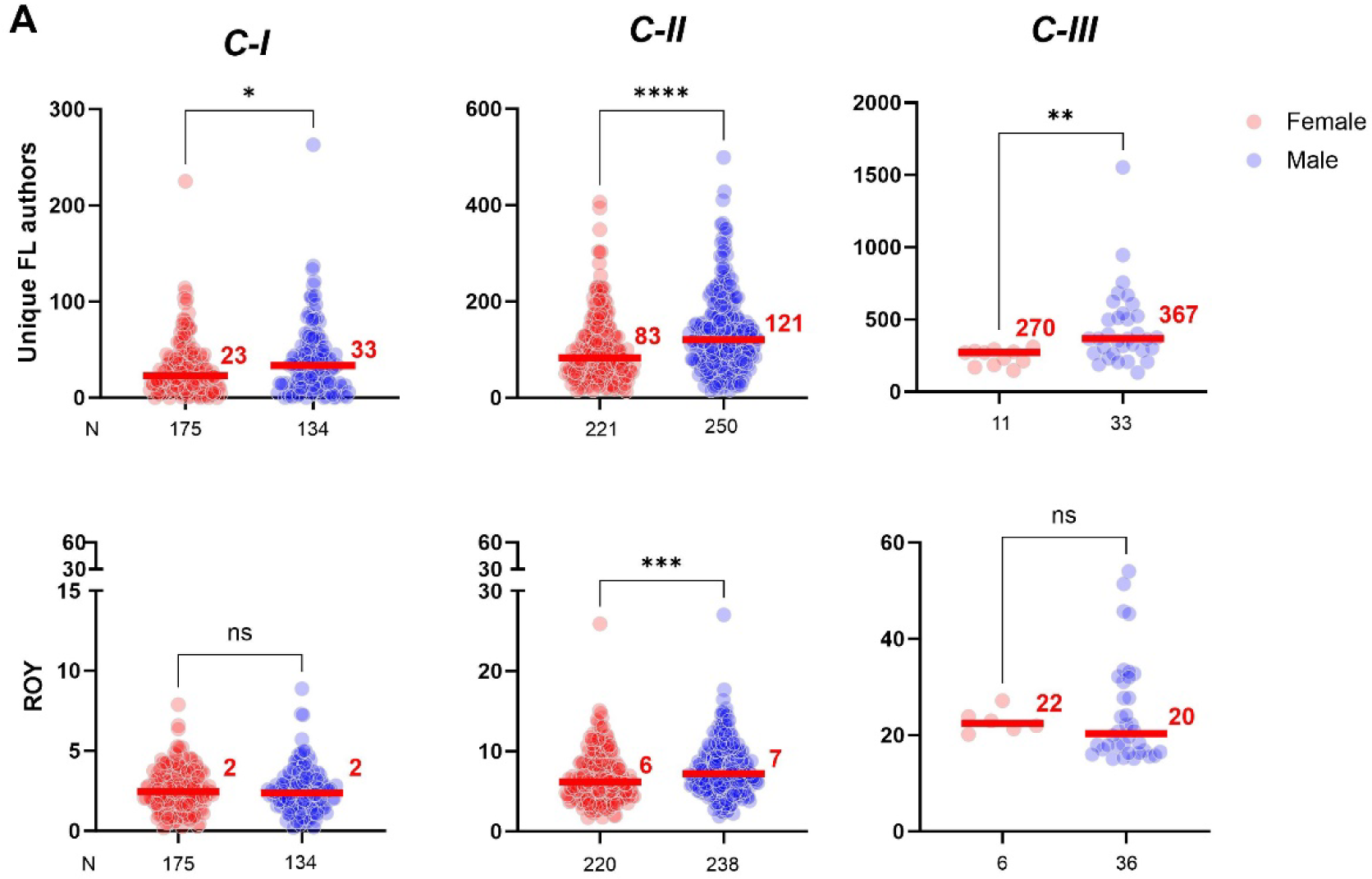
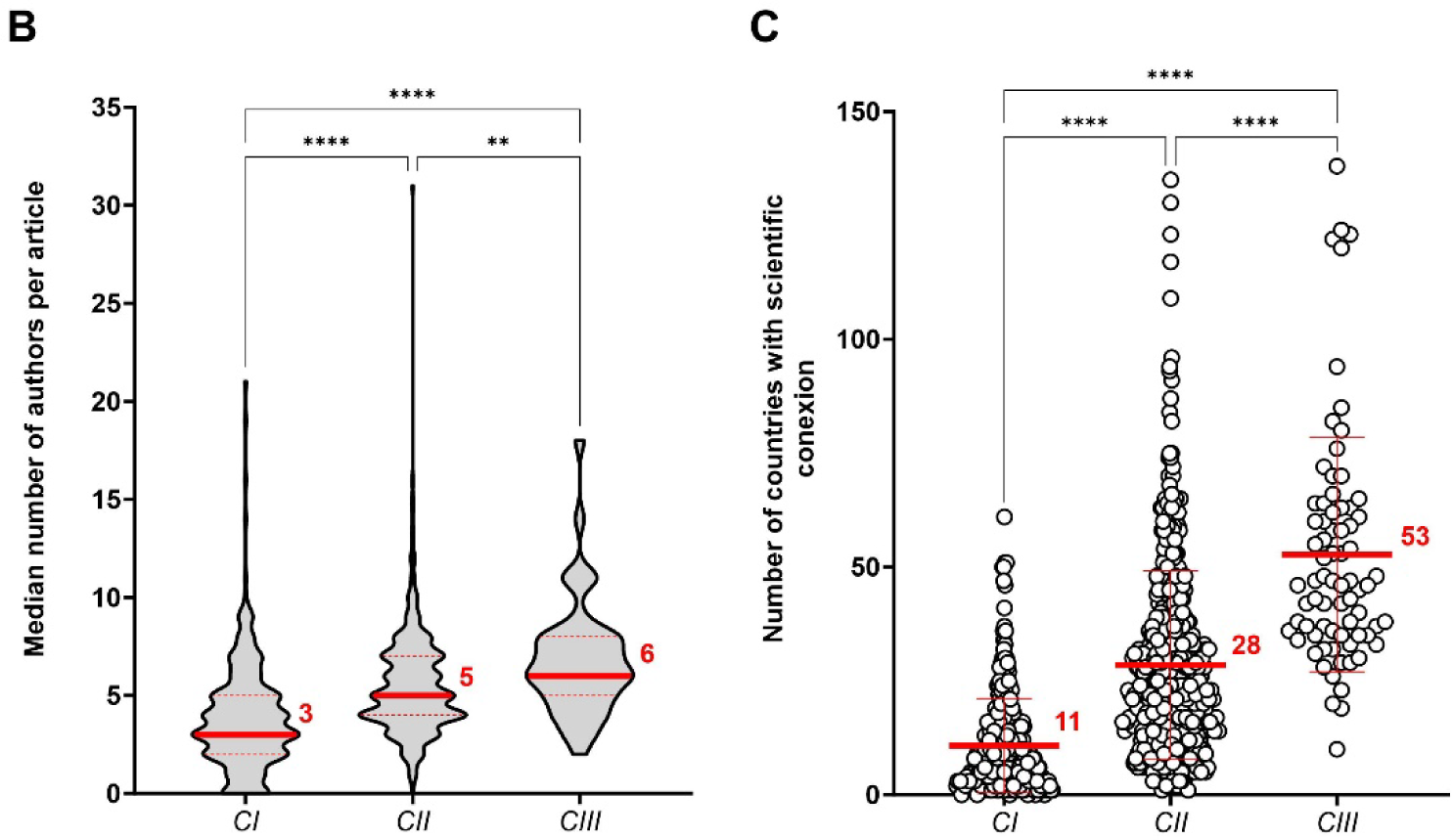
Size-unique differences in the collaborative networks between female and male scientists do not impact their scientific output nor funding. **A)** Peer-comparisons within same *C* level demonstrate that even female scientists tent to have smaller networks than male size (upper panel), this do not play in disadvantage to female productivity (lower panel). B) The generation of knowledge by research teams typically involves around 3 to 6 individuals. C) A scientist’s sphere of influence shows exponential growth as his/her motivational factors to engage in collaborative efforts move up in the *C* level categories.

### Motivational factors for collaborative efforts, and *C* level

Lastly, we set to probe whether *C* score levels can reflect powerful insights on the character and competencies necessary to achieve success in science. Hence, we applied the *C* score level model to our Nobel awardees and ASM-Retraction databases. For the Nobel Laureates (**Fig.9A-C**), as was shown for the tenure-track professors, collaborative efforts demonstrated a highly significant increase as stepping up the *C* levels (Tukey’s multiple comparison test, 40, 205, and 378 unique FL authors, for *C-I*, *C-II*, and *C-III*, respectively). Interestingly, over 60% of *C-I* and *C-II* levels correspond to awardees in the categories of Economics and Physics, and Physiology and Chemistry, respectively, whereas *C-III* level is mostly represented by Physics and no Economics (**Fig. 9A**). The median number of authors ranged from 1 to 5 individuals displaying significant differences only when comparing *C-I* level with *C-II* and *C-III* (Kluskal-Wallis test, 1 vs 4 vs 5, respectively) (**Fig. 9B**). Similarly, the sphere of influences in their scientific world also denoted significant growth as *C* level augmented (Kluskal-Wallis test, 9 vs 28 vs 31, respectively) (**Fig. 9C**). For the ASM-Retraction individuals (**Fig. 9D-F**), perhaps due to the reduced dataset only one individual from the ASM sub-group exhibited a *C* score above 0.8; therefore, subsequent analyses only focused on individuals categorized within *C-I* and *C-II* levels. While the collaborative networks of these two sub-groups tended to increase as moving up in the *C* level categorization, when comparing ASM fellows vs Retraction-related individuals, the number of unique FL authors tended to be smaller for Retraction-related individuals vs ASM fellows, with significant differences for the *C-II* level (Mann Whitney test, 54 vs 35 for *C-I*, and 174 vs 94 for *C-II*) (**Fig. 9D**). Estimation of the median number of authors per article exhibited statistically undistinguishable differences for either *C* level category (Mann Whitney test, 3.5 vs 4.0, and 6.5 vs 5.5, respectively (**Fig. 9E**). Remarkably, significant differences were found for both *C* level categories in their sphere of influence between ASM fellows and Retraction-related individuals (Mann Whitney test, 11 vs 4, and 30 vs 13 countries for *C-I* and *C-II*, respectively) (**Fig. 9F**). Altogether, these results reinforce that small research teams with an average of 5 members tend to be the ones leading breakthroughs in science and technology. In the case of the Nobel Laureates, motivational factors driven academic success do not seem to be strictly related to a research field; while in the sub-group of ASM fellows and Retraction-related individuals further work is needed to elucidate deeper patterns to revel in science. Beyond question, scientists gain visibility with collaborative efforts, boost their productivity, and bolster the ability to expand knowledge influencing their surroundings.

**Figure. 9.**
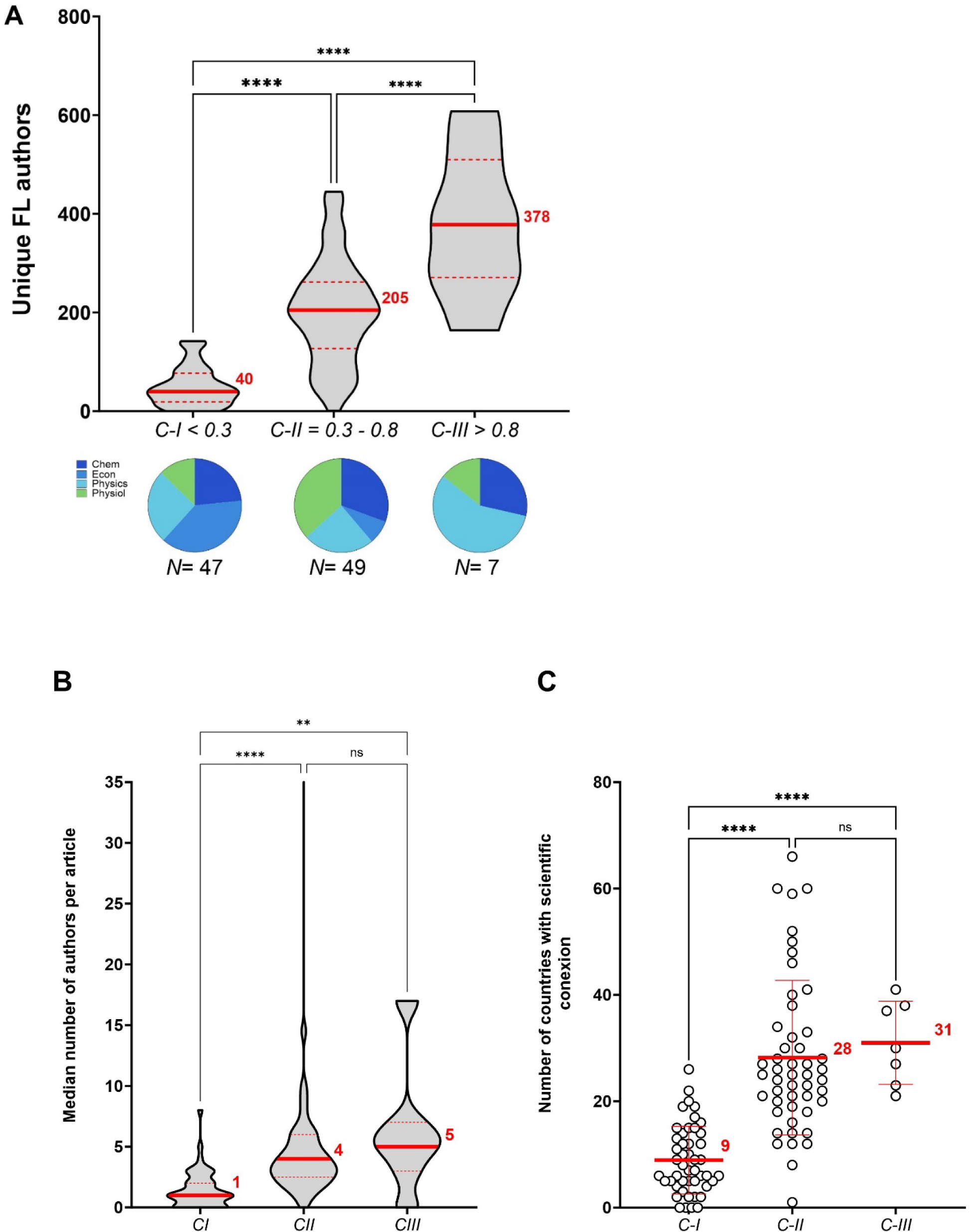
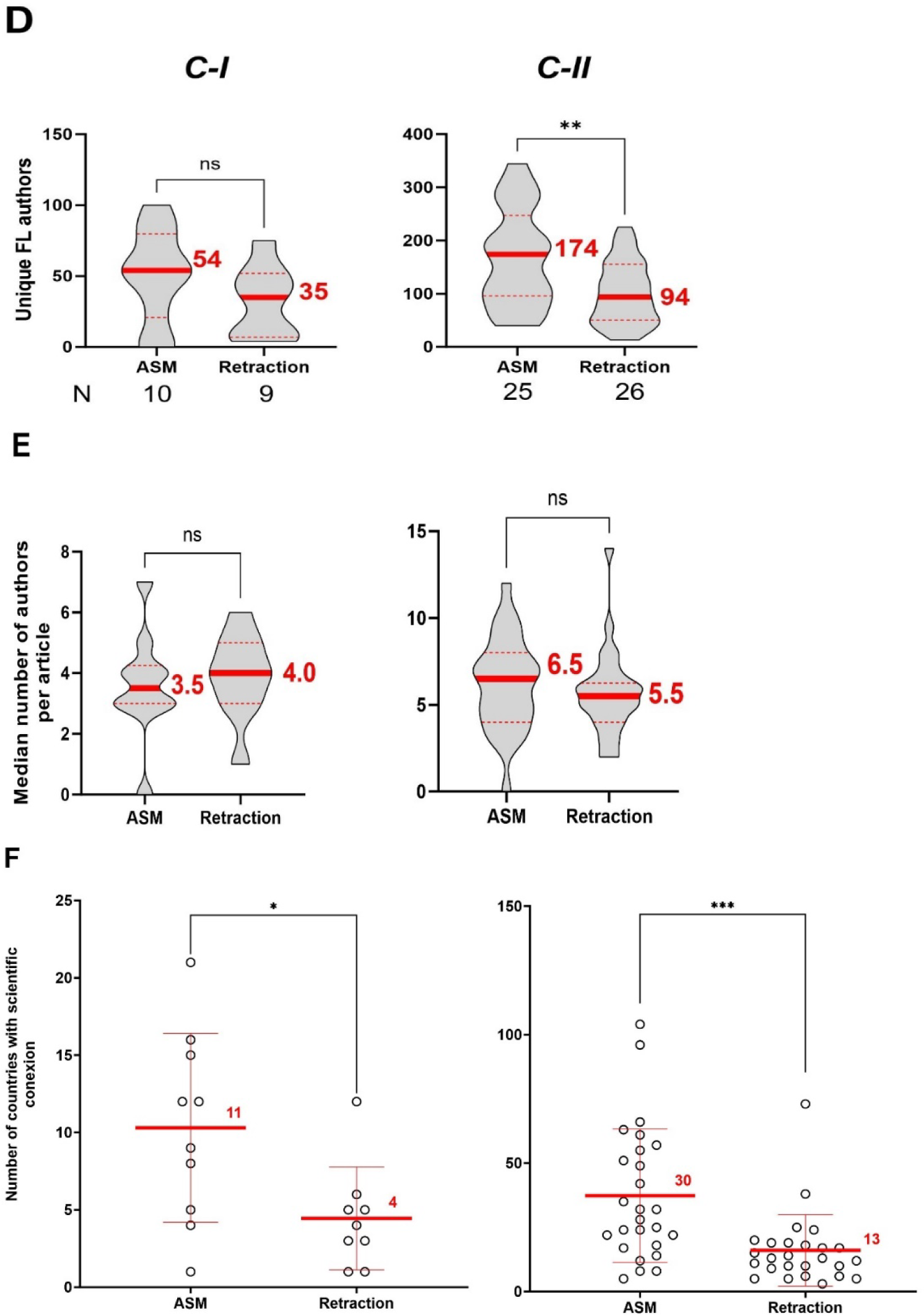
Insights on the characteristics of research collaboration networks of two distinguishing sub-groups based on the *C* score levels. **A)** Size of collaborative networks of Nobel Laureates from 2010-2020 and compositional distribution (pie chart) according to prize categories. **B)** Estimated size of Nobel Laureate’s scientific teams range from 1 to 5 members. **C)** As for academic tenure-track professors, Nobel Laureate’s sphere of influence expanded in proportion to the size of their collaborative networks. **D-F)** Structural and evolutive comparisons between collaborative networks of ASM fellow and Retraction-related scientists according to their *C* level.

## Discussion

In the dynamic landscape of the 21st century, effectively tackling multifaceted societal challenges across health, social, environmental, energy, and technological domains necessitates the pursuit of cross-disciplinary solutions. Collaboration among experts from diverse fields is paramount to provide a holistic perspective and craft comprehensive solutions for these intricate problems. The rapid progression of technologies has ushered in an era of vast data sets and an expanded spectrum of research questions, surpassing the capacity of individual researchers. Acknowledging this reality, research endeavors are increasingly adopting a collaborative approach across virtually all fields, capitalizing on the synergy derived from diverse skills and expertise. Remarkably, around 90% of work in science and engineering disciplines is now accomplished through collaborative efforts [10], emphasizing the pivotal role of scientific teamwork in navigating the intricacies of contemporary research challenges. However, a disconcerting facet of the current scientific landscape lies in the misaligned incentives of its economic system. From promotions to cash bonuses, the prevailing reward system of the scientific enterprise tends to prioritize productivity over scientific rigor, inadvertently creating incentives to cheat emerging from the fear of loss [29, 30]. This unhealthy hypercompetitive culture, driven by scarcity and a winner-takes-all system, risks undermining public confidence in the scientific method [6, 31]. To preserve the integrity of science, it is imperative to reevaluate the economic system of science and shift towards acknowledging the unwritten norms of “scientific civility.”

Competition is an ubiquitous and self-limiting natural process mostly driven by a shortage of resources [32], and science is not exempt of it. As previously argued by Casadevall and Fang [33], perhaps an initial step towards recognizing science as an interconnected and cooperative community is at the level of academic promotions and scientific award committees reviewing an individual’s scientific performance from an integrated perspective, thereby fostering a culture that values collaboration, collegiality, and openness. We propose the *C* score as an index to gauge an individual’s ability to cultivate extensive, sustained, and multicultural collaborations over time, leveraging the strengths and expertise of professionals from different fields. Computed through a simple additive equation based on an individual’s historical record of scholarly productivity and collaborative efforts, the *C* score embodies the human endeavor of science, acknowledging that individuals are uniquely defined by their histories. The score ranges from 0 to 1, one representing a human being as an integrated whole that experiences physiological and psychological aspects.

Regarding linear measurement of the productivity component: A scientist’s productivity is influenced by multiple factors that vary throughout their academic careers, from creativity and ability to publish, to tacit knowledge, social networks, and reputation [34, 35]. The expansion of the scientific literature, measured by the number of articles indexed in the Web of Science database per year, is exponential with an average doubling period of 15 years, while the space of scientific ideas only expands linearly [36]. Furthermore, a period of 15 years for individuals holding a Ph.D. in academia, based on an average of 4-5 years to achieve a milestone within an academic career, also reflects: i) Median length of total time invested during their graduate studies, post-graduate training and then becoming eligible for a tenure position [37]; ii) Development of an independent career; and iii) Significant growth on their sphere of influence. During the length of an academic career, most academic scientists are immersed in teaching and learning with recurring roles as mentee and mentor. Hence, an individual’s productivity was assessed for a maximum 15- year timeframe using 5 categories of productivity level while their collaborative network was quantified into two timeframes: 12-year and a 3-year periods. We focused on full-time faculty members with a tenure status (i.e. Assistant Professor, Associate Professor, and Professor) in three top Schools of Public Health with very high research activity [38]. The 12-year timeframe attempts to assess collaborative efforts with other scientists who may have been Ph.D. and/or post-doctoral supervisors, post-doctoral peers, mentees or contemporary colleagues at early tenure stages among others; while the 3-year timeframe endeavors to measure the development of collaborative projects with current Ph.D. and post-doctoral mentees, previous Ph.D. and post-doctoral mentees who have become independent researchers, and individuals from different areas of specialization with new perspectives to tackle an old problem or solve a new one. In the workplace, productivity refers to how much work (output) is generated over a specific time. Scholarly publications (e.g. research articles, books, conference proceeding, etc.) are the primary mode of communication in science helping disseminate knowledge. In academia, the traditional gold-standard to measure productivity of an academic scientist is the number of publications (Research Output) generated during the length of their career, which contributes to increased knowledge and differs from contributions to scientific progress [36]. Sustained high productivity is rare [39, 40]; the overall process of a scientist’s individual productivity depends on multiple factors, independent hurdles fueling the infamous “publish or perish” dictum. From identifying a good a problem to work on and recognizing worthwhile results, to conveying the relevance of those finding in satisfactory writing, capitalizing on constructive feedback, and showing determination to submit the paper for journal after multiple edits, excel performance in scientific productivity is bounded. In other words, a scientist’s productivity is multiplicative, and it follows is lognormal distribution [39, 41]. Our study with tenure-track professors reproduced those results (**Fig. 1A**) and revealed that this pattern is independent of the career track (**Fig. 1B**) and gender (**Fig 1C**) demonstrating that prolific productivity is motivated by intrinsic factors of each individual, encompassing a set of technical, emotional and psychological skills to successfully juggle the competitive arena of scientific performance.

Regarding the assessment of the collaborative network component: as we delve into our research, each person in our sample group has the potential to contribute significantly to the existing body of knowledge by employing various approaches to address a recognized problem. To illustrate a case, the global effort to eradicate malaria involves a broad spectrum of scientists specializing in different facets of the issue, such as drug discovery, the human host, mosquito host, parasite species, as well as environmental and regulatory policy-making, among others. Each individual in this context acts like a node within a network, capable of acquiring, transmitting, or generating new knowledge. Unlike alternative approaches to quantify collaborative work that use complex algorithms focus on co-authorship or citation analyses, where all co-authorships are treated equally and can be biased towards well-established research and topics, the *C* score provides a holistic view of relationships with a network considering multiples approaches. Our collaborative network quantification assesses the simplest degree centrality measurement for understanding social networks, as the total number of collaborators (node connectivity) and evolution of collaboration over time by capturing the strength of ties (published articles) between collaborators. Furthermore, as scientists work together in cooperative teams, they have the power to revolutionize science and technology with innovative ideas or advance existing ones by drawing on the diverse scientific backgrounds and perspectives within the group. The development and expansion of these collaborative networks depend on a scientist’s values, personal aspirations, and needs over time. That being so and within the “scientific civility” framework, the *C* score also provides insights into the breadth of collaboration accounting for global collaborative network when scientific interactions involved individuals from 4 or more continents. This approach serves as a diversity metric that favors chances for the existence of collaborative projects with truly diverse researchers affiliated with institutions from distinct cultural backgrounds.

Some caveats in our analysis were noted. Our *C* score calculation between the academic ranks analyzed in this study was not adjusted to the appointment year since that information is not publicly available **(Figs. 4A-B)**. We anticipate that accounting for it might provide a clearer picture of an individual’s current networks activity. For the estimation of academic career length, we used the very first paper published, which may not reflect the appropriate starting point if it represents a small contribution in a field with which one is no longer involved (**Fig. 4A**). The *C* score distribution in social sciences such as Economics tends to the lower end mainly attributed to a low ROY in the form of publications, which could be possible corrected by taking in consideration a more adequate scholarly output per year for this field such as books (**Fig. 5A**). The *C* score also cannot predict whether collaborative networks are forged in response to selfish and malicious competition or cooperative and joyful interactions between investigators (**Figs. 6A-C**); nor fairly allocate fractional credit share in multi-authors papers. While total number of collaborators and unique FL authors strongly correlates with the *C* score (**Figs. 7A-B**), this metric cannot project the *C* score of an individual at a future time since it is not cumulative; both scholarly productivity and collaborative network growth require active and regular input. Boosting scholarly productivity by means of “salami slicing” cannot be discriminated against with the *C* score [42, 43]. Finally, the importance or significance of the publications (i.e., number of citations) used in calculating the *C* score is not considered in the calculation. Our metric did not compute journal impact factors.

Despite the foregoing limitations, normalizing productivity on a yearly basis enables the comparison of scientists across different ages who hold the same academic ranks. Within the academic setting, the *C* score serves as a valuable tool for assessing the upward trajectory of a scientist’s career from Assistant Professor to Professor (**Fig. 4B**). This aspect is crucial for mentoring and developing human talent. For instance, if an individual’s *C* score falls below the expected range for their cohort during professorial rank progression, it can signal the need for intervention to ensure continued academic success through additional mentoring and advice. The assessment of collaborative networks concerning space and time allows for tracking and quantifying multicultural, sustained, and unique collaborative efforts in two distinct timeframes of a scientist’s career. This is an essential characteristic of inclusive and diverse cooperative work, significantly impacting the workplace environment. Moreover, mindful of the limitations of assigning binary genders, the *C* score reveals statistically undistinguishable differences between female and male tenure-track professors of equal academic rank (**Fig. 4D**). In contrast to the *h* index, which captures a passive cumulative impact of an author’s scholarly performance [12], the *C* score exhibits a dynamic behavior (**Fig. 5C**). It can decrease, increase, or remain unchanged over time, responding to levels of engagement with collaborative projects. This reflects the cumulative relevance and productivity of an individual at a defined point of their academic career. Thus, a high *C* score can be assumed as an indicator of an accomplished and collaborative scientist, but the opposite is not necessarily correct. Calculating and averaging the *C* score annually can provide a better indicator of consistencies in “scientific civility” over the lifetime of a scientist (**Fig. 5C**). This indicator assesses academic performance in terms of a scientist’s active collaborative networks that prevail over institutionally limited economic resources and avoids inequalities related to the length of an individual’s academic career, field of investigation, and gender. The *C* score parameter also reflects the likelihood that an individual will be highly recognized by their scientific achievements and that comparisons in productivity between men and women can be avoided.

Lastly, as outlined by Abraham Maslow, human motivation revolves around the pursuit of fulfillment and personal growth [27, 28]. Life experiences can significantly influence an individual’s needs, either enhancing or disrupting their progress-this holds true for academic career tenure-track advancement. Aligning with the observed progressive pattern of median *C* scores during the tenure-track career, we propose that academic performance is driven by a three-level model of motivational development based on physiological and psychological needs. The model integrates productivity and collaborative efforts, defining three stages for professional growth of tenure-track professors in academic institutions from *C-I* to *C-III*. *C-I* represents deficiency-motivated individuals, primarily driven by physiological, safety, and social needs such as securing food and housing, establishing an independent line of investigation, and gaining peer recognition. *C-II* includes scientists motivated by a deeper appreciation of their surroundings, finding satisfaction in life’s order, elegance, and harmony. *C-III* encompasses those who strive for their highest potential or assist others in achieving it, motivated by values transcending personal self and rooted in a profound sense of belonging within the biosphere (**Fig. 7C**). Therefore, framing our data within the *C* level of motivational development unveils significant factors influencing academic performance. Notably, the *C* score level shows gender-specific and statistically significant differences in the size of collaborative networks. As *C* levels ascend the hierarchy, both female and male scientists exhibit an expansion in the size of their collaborative patterns, emphasizing the importance of trustful relations are a key element for scientific success (**Fig. 8A, upper panel**). It is noteworthy that, irrespective of the *C* level, networks of female scientists were consistently smaller than those of their male counterparts. This finding aligns with previous studies in the field of computer science, where female networks are characterized by collecting knowledge in tightly knit communities while men tend to seek innovations across structural holes [44]. Conversely, our study suggests that this structural difference in their networks do not impede female compared to males. In agreement with previous studies on the structure of research teams, we found that size of collaborative research teams, defined by the median number of authors per article, seem to be structured by a core group of scientists, typically ranging from 3-6 members, highly interconnected to promote effective communication without jeopardizing creativity or productivity [26]. This pattern was gender-independent but driven by similar motivational factors.

Furthermore, application of the *C* level categorization to the Nobel Laureates highlights productivity and social conventions patterns characteristics on particular scientific fields such as Economics and Physics, in which solo-authored work mostly in the forms of books or the publication of research articles with a large number of co-authors are typical, respectively. In these cases, one could think that a possible way to correct the *C* score would be adapting the scholarly output for the most significant output of the field whether it is research-, teaching-, or practice-related. On a different angle, notable differences on the structure and evolution the collaborative networks between ASM fellow and Retraction-related individuals were highlighted by this study; further analysis are in progress to gain deeper understanding of these dynamics aiming to find solutions that can channel competition and foster beneficial cooperative efforts to enhance the scientific enterprise. In this regard, multiple metrics have been proposed to account for the collaborative effect [45, 46]: fractional credit share in muti-authored paper, distinguishing amongst roles played by each co-authors such as first and last authorships; to identify scientific leadership/eminence. No single metric can summarize the myriad contributions scientists make to the scientific enterprise and society, including but not limited to discoveries, citations, teaching, mentoring, developing academic curricula, reviewing, organizing scientific meetings, serving on ethical committee, and editorial boards. Nonetheless, the *C* score may reflect the ability of scientists to build and sustain dynamic working groups based on trust, consistency, and clear communication. Individuals with higher *C* scores are perhaps more likely to think about team science [7], and possess strong communication skills that promote competition while containing conflict to bolster the emergence of creative and multicultural cooperative solutions. These components of scientific productivity, generativity, and leadership require a strong foundation of civility.

In conclusion, we propose the *C* score as a new research indicator for evaluating the academic performance of an individual in terms of scholarly output and collaborative efforts at a certain timepoint of their career. The integral perspective of assessing academic performance on sustained productivity and collaborative contributions over time rather than on a single metric (e.g. number of articles, number of citations, *h*-index, etc.) may shield from manipulation and to become vulnerable to Goodhart’s law: “When a measure becomes a target, it ceases to be a good measure”[47]. For administrators and other professionals in decision-making scenarios, the *C* score, along with other indicators such as the *h* index, would support a more integrative and holistic assessment of an individual’s academic performance. By championing a reward system aligned with the principles of “scientific civility,” we can pave the way for a more robust and ethically sound scientific community.

## Materials and Methods

### Sample

The analysis was conducted from 2006 to 2020 with a total of 1000 academic individuals (825 Tenure-Track Professors, 103 Nobel Laureates from 2010-2020 in four distinct categories, 37 ASM Fellows, and 35 USA scientists involved in retracted articles. The study relied on analysis of publicly available bibliometric data and no interactions of any kind (e.g., face-to-face, on paper, or in electronic realms) with any individual was conducted. Nonetheless, the study was submitted to The Johns Hopkins Medicine Institutional Review Boards (JHM IRBs) for reviewing and qualified as exempt research under the U.S. Department of Health and Human Services (DHHS) regulations.

### Data

To collect the information used for this study, we used the SCOPUS database curated by Elsevier. We performed SCOPUS author, document, and affiliate searches to retrieve the entire body of scholarly literature associated with each individual via unique Scopus ID. This data was collected through the SCOPUS APIs using the Elsapy Python package (version 0.5.0). Collaborator affiliations for calculating global network scores are based on the affiliation of each author at the time of publication for each given article. Data used for this study were downloaded between 2021-2022 and included the full years between 2006 and 2020. Data was available from The Center for Scientific Integrity, the parent nonprofit organization of Retraction Watch, subject to a standard data use agreement. Gender was assumed by searching for images of an individual’s name using the Google search engine. For one individual, we were not able to assign a gender. All the primary information used in this study was obtained from publicly available databases, such as university faculty directories, Nobel Prize winner lists, etc. Quality of the code was assessed with a sample population (around 15%) by accessing the SCOPUS database using an individual’s full name and manually measuring all parameters associated to the ROY and CN variables.

### Statistical analysis

Relationships between the *C* score and *h* index were calculated using a Spearman correlation coefficient between the two values for each given dataset; 95% confidence intervals were used to summarize association. Correlations were not calculated for any grouping that included < 4 individuals. Analyses were performed in GraphPad Prism (version 10.0.1).

## Acknowledgments

We thank Gayane Yenokyan at the Institute for Clinical and Translational Research for her insight and assistance in the biostatistical analysis, and Marie J. Hardwick, Radames J.B. Cordero, Marcio Rodrigues, and Gundula Bosch for helpful conversations. The illustration in Figure 2 was made using Biorender.com.

---

**Supp. Figure 1.**
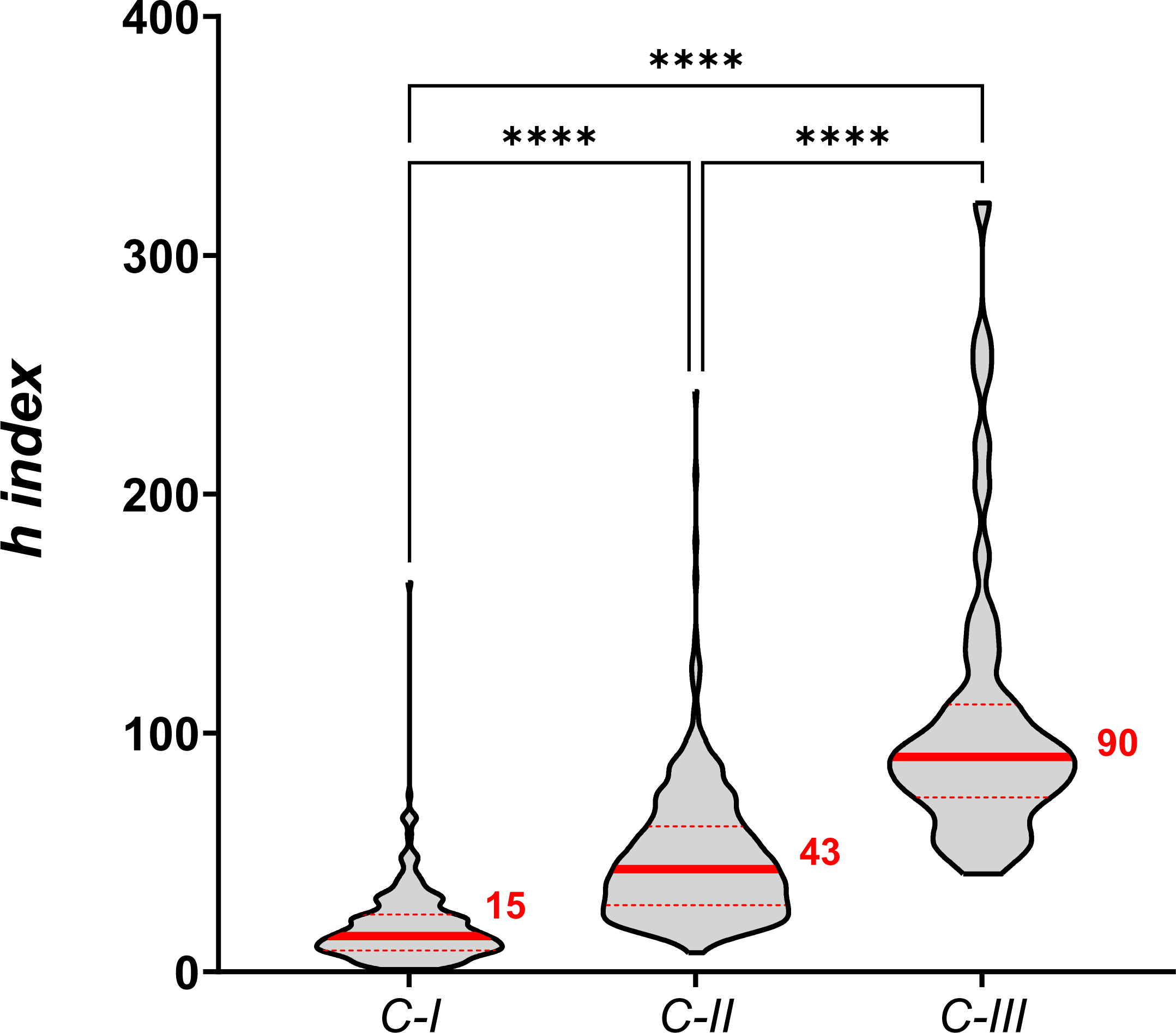

