## Supplementary figures and images for "Scientific civility and academic performance"

### Supp Fig1

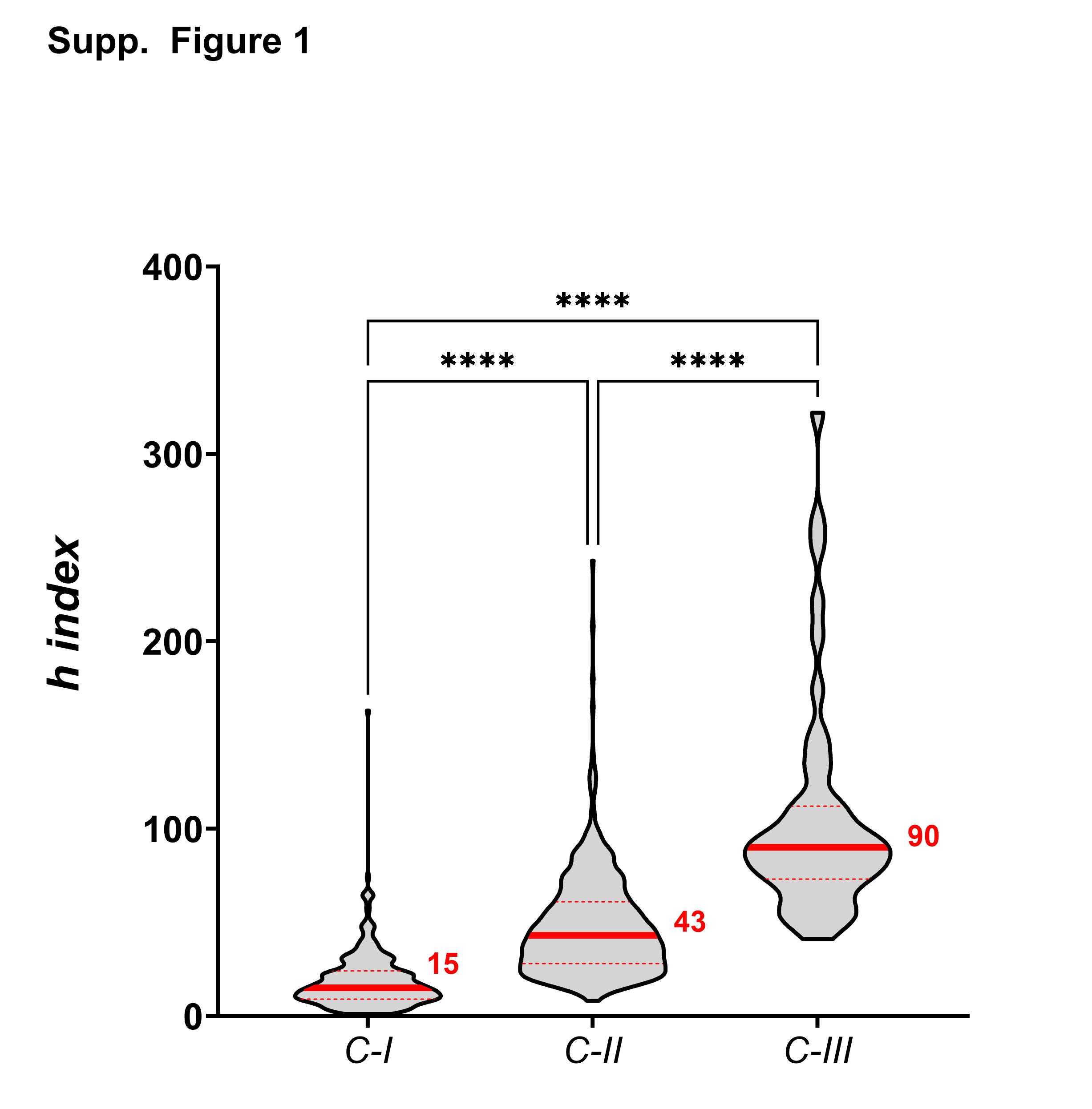
